# Dosage-sensitivity shapes how genes transcriptionally respond to allopolyploidy and homoeologous exchange in resynthesized Brassica napus

**DOI:** 10.1101/2021.11.16.468838

**Authors:** Kevin A. Bird, J. Chris Pires, Robert VanBuren, Zhiyong Xiong, Patrick P. Edger

## Abstract

The Gene Balance Hypothesis (GBH) proposes that selection acts on the dosage (i.e. copy number) of genes within dosage-sensitive portions of networks, pathways, and protein complexes to maintain balanced stoichiometry of interacting proteins, because perturbations to stoichiometric balance can result in reduced fitness. This selection has been called dosage balance selection. Dosage balance selection is also hypothesized to constrain expression responses to dosage changes, making dosage-sensitive genes (those encoding members of interacting proteins) experience more similar expression changes. In allopolyploids, where whole-genome duplication involves hybridization of diverged lineages, organisms often experience homoeologous exchanges (HEs) that recombine, duplicate, and delete homoeologous regions of the genome and alter the expression of homoeologous gene pairs. Although the GBH makes predictions about the expression response to HEs, they have not been empirically tested. We used genomic and transcriptomic data from six resynthesized, isogenic *Brassica napus* lines over ten generations to identify HEs, analyzed expression responses, and tested for patterns of genomic imbalance. Groups of dosage-sensitive genes had less variable expression responses to HEs than dosage-insensitive genes, a sign that their relative dosage is constrained. This difference was absent for homoeologous pairs whose expression was biased toward the BnA subgenome. Finally, the expression response to HEs was more variable than the response to WGD, suggesting HEs create genomic imbalance. These findings expand our knowledge of the impact of dosage balance selection on genome evolution and potentially connect patterns in polyploid genomes over time; from homoeolog expression bias to duplicate gene retention.

## Introduction

The observation of genomic imbalance (Box 1), that changing the dosage of chromosomes or chromosome segments has a more detrimental phenotypic effect than changes to whole chromosome sets (e.g. whole-genome duplication, WGD), led to the appreciation for gene dosage changes as a powerful and important driver of gene expression abundance, quantitative trait variation, and the evolution of genomes (Birchler and Veitia 2007, 2010, 2012). These dosage changes can be divided into absolute dosage changes, related to the direct increased copy number of genes and abundance of gene product, and relative dosage changes, related to the relative copy number of genes that encode proteins connected within networks, pathways, or protein complexes and the resulting stoichiometry of their gene products (Bekaert et al. 2011; Conant et al. 2014). The finding that altered stoichiometry of gene products can have large phenotypic impacts and be highly deleterious for certain classes of genes, especially those involved in highly connected regulatory networks and multimeric protein complexes led to the formulation of Gene Balance Hypothesis to explain the basis of genomic imbalance (Birchler and Newton, 1981; Birchler et al., 2001; Makino and McLysaght, 2010; Birchler and Veitia, 2012). The core of the Gene Balance Hypothesis proposes that changing the stoichiometry of members of networks, pathways, and protein complexes affects the kinetics, assembly, and function of the whole, which incurs negative fitness consequences (Birchler et al., 2005; Birchler and Veitia, 2007, 2010, 2012). A valuable contribution of the Gene Balance Hypothesis, and the study of relative gene dosage, is that it provides a theoretical framework to connect observations of genetic and genomic variation in the present to the past and to predictions of future evolutionary trajectories of genes and gene networks. Dosage balance selection, the selection on relative dosage to maintain the stoichiometric balance of gene products in the face of changes in gene dosage, has since been shown to influence genome evolution in important and predictable ways (Conant et al. 2014). This includes expectations of karyotype evolution (Xiong et al. 2011), duplicate gene retention and loss (Edger and Pires 2009, Makino and McLysaght, 2010; De Smet et al. 2013; Tasdighian et al. 2017; Emery et al. 2018, Gout et al. 2023), gene expression (Coate et al. 2016; Shi et al. 2015; Hou et al. 2018; Song et al. 2020; Shi et al. 2021; Yang et al. 2021), and even potential mechanisms for polyploid formation (Cao et al. 2023).

#### Box 1.

Glossary

Box 1. Glossary
|  |  |
| --- | --- |
| Absolute Dosage | The direct copy number of genes – absolute dosage will not necessarily track the abundance of gene product. |
| Allopolyploidy | Polyploidy from the hybridization of genetically diverged individuals. |
| Autopolyploidy | Polyploidy from the hybridization of genetically similar individuals or direct duplication of an individual's genome. |
| Dosage-Balance Selection | The selection to maintain the stoichiometric relations of the products of dosage-sensitive genes. |
| Dosage-Sensitive Genes | Genes for which a change in copy number alters expression and protein abundance and interrupts the stoichiometric balance of their gene products with those from other genes – in this study dosage-sensitivity is inferred based on patterns of duplicate gene retention. |
| Gene Balance Hypothesis | The hypothesis that the stoichiometry of members of multisubunit complexes can affect the amount of functional complete product, which in turn, affects patterns of gene expression, and ultimately, the phenotype and evolutionary fitness |
| Genomic Imbalance | The more severe phenotypic effects of altering the number of a single chromosome or chromosome segment compared with altering the number of all chromosomes. |
| Homoeologs | Orthologous genes or chromosomes that diverged in the polyploid progenitors and were reunited by polyploidization. |
| Homoeologous Exchange | Recombination between homoeologous chromosomes that swaps genomic regions between subgenomes and can result in deletions, duplications and translocations of homoeologous regions. |
| Homoeologous Exchange Response Variance (HERV) | The coefficient of variation (SD/Mean) of the foldchange between expression in an HE region and the balanced (2:2) expression for that region (summed progenitor expression) for gene pairs affected by an HEs & within a GO term. |
| Homologs | Genes or chromosomes that share a common origin. |
| Orthologs | Genes or chromosomes from different species that arose by a speciation event. |
| Paralogs | Genes that arose by a duplication event. |
| Polyploidy | Having two or more complete sets of chromosomes. |
| Polyploid Response Variance (PRV) | The coefficient of variation (SD/Mean) of the foldchange between tetraploid expression and the midparent expression for all gene pairs from balanced (2:2) genomic regions & within a GO term. |
| Relative Dosage | The relative copy number of genes that encode proteins connected within networks, pathways, or protein complexes-- relative dosage is primarily relevant for dosage-sensitive genes. |
| Subgenome | A complete set of chromosomes derived from a progenitor species in an allopolyploid. |

Comparative genomic studies have supported predictions from the Gene Balance Hypothesis, showing that for certain classes of genes, duplicate gene retention shows biased patterns depending on whether a gene is duplicated by whole-genome duplication or by small-scale duplications. This pattern of biased gene loss is best explained by dosage balance selection which selects against imbalanced stoichiometry of dosage-sensitive genes and constrains their range of possible dosage. Duplicate copies from many transcription factor families, genes involved in signaling pathways and multimeric protein complexes, and others tend to be retained more than expected after whole-genome duplication, which maintains the relative dosage of these genes, and duplicates from small-scale duplications, which perturb relative dosage, tend to be retained less than expected (Blanc and Wolfe, 2004; Maere, 2005; Paterson et al. 2006; Thomas and Freeling, 2006; Freeling, 2009; Edger and Pires, 2009; De Smet et al., 2013; Conant et al., 2014; Li et al., 2016; Tasdighian et al., 2018). This pattern of preferential retention from whole-genome duplication and loss from small-scale duplication has been called “reciprocal retention” (Freeling, 2009; Tasdighian et al., 2018). Many of these studies have focused on ancient WGD events, where genomes have returned to a diploid-like state, leaving the immediate transcriptional impact of large-scale gene dosage changes less well understood.

Several authors have recently investigated the expression responses caused by aneuploidy and polyploidy (Coate et al. 2016; Hou et al. 2018; Song et al. 2020; Shi et al. 2021; Yang et al. 2021). Coate et al. (2016) and Song et al. (2020), in particular, attempt to connect observed patterns of long-term duplicate gene retention to short-term duplicate gene expression responses. They use tenets of the Gene Balance Hypothesis to predict two patterns in short-term expression response. First, genes that are reciprocally retained after whole-genome duplication (e.g. those that are highly connected in gene networks, involved in multicomponent protein complexes, etc.) should experience a change in gene expression in response to genome duplication so that selection is able to act on dosage changes. Second, expression changes should be less variable for all genes in a network or functional class, what they call a “coordinated response”. Coate et al. (2016) address this question using natural soybean (*Glycine* L.) allopolyploids with an origin ∼500,000 years ago and known diploid progenitors, while Song et al. (2020) use three *Arabidopsis thaliana* autopolyploid/diploid pairs. Both studies determined that genes in reciprocally retained GO terms showed a less variable expression response to polyploidy than genes from non-reciprocally retained GO terms, suggesting that genes under selection to maintain genomic balance have a constrained transcriptional response to dosage changes (Coate et al. 2016; Song et al. 2020). In the case of Song et al. (2020), the use of synthetic polyploids revealed that this expression response is an immediate response to altered gene dosage. Differences in expression modulation of different functional classes were also observed in synthetic polyploid and aneuploid lines of *Arabidopsis* (Hou et al. 2018) and maize (Shi et al. 2021). The use of autopolyploids or aneuploids facilitates the isolation of different types of dosage changes for investigations of genomic balance but misses features unique to allopolyploids, which involve the hybridization of evolutionary diverged lineages and the doubling of genomic material, like homoeologous exchange or subgenome expression bias that may affect the maintenance of relative gene dosage and balanced stoichiometry.

Early studies in resynthesized allopolyploids showed extensive genetic changes in a short period of time (Song et al. 1995). Subsequent investigations showed major genome structural changes from the first meiosis after polyploid formation, primarily in the form of homoeologous exchanges in which recombination among homoeologous regions results in the partial or complete deletion and duplication of chromosomal segments (See Deb et al. 2023 for a recent review, see also Sharpe et al. 2005; Osborn et al. 2003; Jenczewski et al. 2003; Pires et al. 2004; Gaeta et al. 2007; Nicolas et al. 2007; 2012; Szadowski et al. 2010; Xiong et al. 2011; Chalhoub et al. 2014; He et al. 2017; Rousseau-Geutin et al. 2017; Samans et al. 2017; Stein et al. 2017; Hurgobin et al. 2018; Lloyd et al. 2018; Pele et al. 2018; Mason and Wendel 2020; Bayer et al. 2021; Chawla et al. 2021; Ferreira de Carvalho et al. 2021; Higgins et al. 2021; Xiong et al. 2021; Orantes-Bonilla et al. 2022; Cao et al. 2023). These rearrangements continue to accumulate over time, generating genomic diversity in early polyploids (Gaeta and Pires 2010; Xiong et al. 2011; Mason and Wendel, 2020). Homoeologous exchanges are often destructive to the organism and meiotic stability is more frequently observed in natural polyploids compared to resynthesized and it is likely that meiotic stability is under strong selection in natural polyploid populations (Gaeta and Pires, 2010; Rousseau-Gueutin et al. 2017; Pele et al. 2018; Xiong et al. 2020; Gonzalo et al. 2019; Gaebelein et al. 2019; Ferreira de Carvalho et al. 2021). At the same time, homoeologous exchanges generate phenotypic novelty in resynthesized polyploids (Pires et al. 2004; Gaeta et al. 2007; Wu et al. 2021) and are frequently observed in natural polyploids (Chalhoub et al. 2014; Lloyd et al. 2018; Edger et al. 2019; He et al. 2017, Chawla et al. 2021). Homoeologous exchanges may be under genetic control (e.g., Jenczewski et al. 2003; Higgins et al. 2021), may affect meiotic stability (Xiong et al. 2021), may underlie gene presence-absence variation and agronomically valuable quantitative trait loci in *Brassica napus* (Samans et al. 2017; Stein et al. 2017; Hurgobin et al. 2018; Bayer et al. 2021), and may generate novel, chimeric transcripts as recently observed in several polyploid species including wheat, *Brassica napus, Arabidopsis suecica*, banana, peanut, and synthetic tetraploid rice (Zhang et al, 2020). Less is known about the evolution of homoeologous exchanges.

Additionally, allopolyploid genomes must accommodate inherited and novel expression differences in homoeologous genes which often results in subgenome dominance, where expression is biased in favor of homoeologs from one progenitor genome over others. (Alger et al. 2021; Bird et al. 2018,2021; Wendel et al. 2018). This effect is driven by the merger of evolutionarily diverged genomes, which frequently results in remodeling of epigenetic markers (Madlung et al., 2001; Edger et al., 2017; Bird et al., 2021), alterations in gene regulation (Chen, 2007), and activation of transposable elements (Vicient and Casacuberta, 2012). Importantly, there is also a continuum of polyploidy, as parental genomes within and among species can vary in evolutionary distance, and subsequent genome evolution blurs the distinction between autopolyploidy and allopolyploidy (Stebbins 1947; Leal-Bertioli et al. 2018; Mason and Wendel 2020; Blischak et al. 2023; Bomblies 2023). Subgenome expression dominance has been defined in terms of a subgenome possessing a greater amount of dominantly expressed homoeologs and has been identified in many allopolyploid species, including maize (Schnable et al. 2011) *Mimulus peregrinus* (Edger et al. 2017), garden strawberry (Edger et al. 2019), *Brassica rapa* (Cheng et al. 2012; Cheng et al. 2016), and a population of resynthesized *Brassica napus* (Bird et al. 2021).

Unlike aneuploidy and polyploidy, the impact of gene expression changes from homoeologous exchanges or biased homoeolog expression on dosage balance is largely unexplored. There are reasons to believe homoeologous exchange can alter the balance of gene products in ways that entail specific evolutionary predictions from the Gene Balance Hypothesis. Lloyd et al. (2018) investigated the effect of homoeologous exchanges on expression in natural *Brassica napus* and found that Lloyd et al. (2018) found when homoeologous gene pairs have unequal expression, altering the ratio of homoeologous copies by homooeologous exchange result in dosage-dependent expression changes (i.e. proportional to the change in gene copy number). These expression changes did not accurately compensate to maintain the same level of combined homoeolog expression. Similar results have since been observed in tetraploid wheat lines (Zhang et al. 2022). This expression modulation from HE dosage changes resembles the protein modulation seen in aneuploid and polyploid maize lines (Birchler and Newton, 1981). Therefore, in the presence of unequal homoeolog expression, dosage changes from HEs will alter expression levels of homoeologous gene pairs and, potentially, the stoichiometry of interacting gene products. Since HEs only affect a subset of the genome, the Gene Balance Hypothesis predicts that this change in expression would lead to greater genomic imbalance compared to polyploidy. The GBH also predicts that the constraint on relative gene dosage from dosage balance selection would result in a more similar expression response for groups of dosage-sensitive genes affected by an HE. Whether these predicted patterns hold has not yet been explored.

We analyzed paired WGS and RNASeq data for six independently resynthesized and isogenic *Brassica napus* (CCAA) lines, which are known to accumulate large amounts of genomic rearrangement (Xiong et al. 2011), at three generations to identify homoeologous exchange events that resulted in altered relative dosage of genes and tested two predictions from the Gene Balance Hypothesis regarding the transcriptional response to homoeologous exchanges. The first was dosage-sensitive genes will have a less variable expression response to homoeologous exchange. The second was that the expression response to homoeologous exchanges will be more variable than the expression response to whole-genome duplication.

Based on previous results indicating subgenome dominance in this population of resynthesized lines favoring the BnC subgenome (Bird et al. 2021), we further tested the expression response to homoeologous exchange and whole-genome duplication to see if the transcriptional response to dosage changes differed based on which homoeolog was more highly expressed. Such results may suggest that dosage balance selection can differ between gene pairs due to the direction of subgenome-biased expression. Additionally, individuals from the first, fifth, and tenth generations were examined to see if expression responses changed over time. Our findings provide new understanding of how selection to maintain balanced stoichiometry of gene products affects gene expression and genome evolution across various modes of gene dosage changes in newly formed polyploids.

## Methods

### Sequencing data

We downloaded the whole genome sequences (WGS) and RNAseq data and files for previously identified genomic rearrangements and transcript quantification from leaf samples from Bird et al. (2021) at the associated Data Dryad repository https://doi.org/10.5061/dryad.h18931zjr. These previous analyses identified 26,111 syntenic ortholog pairs between the progenitor genomes, treated as homoeologous pairs from here on.

### Identification of homoeologous exchange events

We used the dosage assignments from Bird et al. (2021). Briefly, read depth ratio of WGS resequencing data for homoeologous pairs were calculated over a 50 gene sliding window with stepsize of one. Homoeologous pairs were assigned 0:4, 1:3, 2:2 3:1, 4:0 based on distorted ratios of WGS reads mapping to one homoeolog over the other along a sliding window of 170 genes with a step size of one gene. To account for uncertainty in alignment and potential cross-mapping, the read depth was split into equal-sized quintiles to assign a dosage (0-20% as 0:4, 20-40% as 1:3, etc). Additionally, a region was only assigned as a homoeologous exchange if 10 or more consecutive genes had WGS read depths within the defined quintiles.

### Expression Quantification

Read count files for these samples had previously been filtered to remove lowly expressed pairs by removing gene pairs with summed TPM < 10, allowing for the potential that one copy is truly silenced. The number of homoeologous gene pairs with expression quantification in these samples ranged from 11,355 to 12,939, while the number of gene pairs affected by genomic rearrangements with expression quantification ranged from 148 for one plant in the first generation to 4606 for a plant in the tenth generation.

### Dosage-sensitivity assignment

To leverage the well-curated gene annotations of *Arabidopsis thaliana*, and the close phylogenetic relationship between *A. thaliana* and the *Brassica* genus, we assigned our *Brassica* gene pairs to the GO category of their *A. thaliana* ortholog. Orthologs between *A. thaliana* and *Brassica oleracea* were identified with Synmap (Lyons et al. 2008) on CoGe (Lyons and Freeling, 2008) and the *A. thaliana* GO annotations were directly assigned to the *B. oleracea* orthologs and from *B. oleracea* to the *B. rapa* syntelogs. Next, we used the GO term dosage response assignments (dosage-insensitive and dosage-sensitive) from Song et al.’s (2020) analysis of gene retention patterns of *A. thaliana* genes to classify our syntenic homoeologs as belonging to dosage-sensitive and dosage-insensitive GO terms. Arabidopsis genes, their associated GO terms and classification from Song et al. (2022), and the identified *Brassica oleracea* orthologs can be found in the supplementary dataset.

### Homoeologous exchange response variance

We included only syntenic homoeolog pairs that diverged from 2:2 dosage ratio (e.g. gene pairs with read-depth ratio less than 0.4 or greater than 0.6), as identified by Bird et al. (2021), to investigate the effects of gene dosage changes. Previous cytogenetic analysis of these lines revealed substantial aneuploidy and partial chromosomal duplication/deletion, especially among the most syntenic chromosome pairs like A1/C1, A2/C2, and C9/A10 (Xiong et al. 2011). To eliminate confounding effects of these kinds of rearrangements, we checked our lines for regions of skewed read depth that covered the majority or entirety of a chromosome.

We fully excluded chromosomes where the majority of plants showed these large regions of skewed read-depth ratios and individually removed cases where large skewed ratios were seen for chromosomes in one sample. Plots of read depth ratios along the genome for each line and generation are shown in Figs S1-S6. This resulted in the removal of syntenic homoeologs from chromosomes A1/C1, A2/C2, and C9/A10 from all lines, and chromosome C4 only for line EL-1100 at generation 10. Parental expression was taken from Bird et al. (2021), where RNAseq from independent libraries of B. rapa acc. IMB218DH and B. oleracea acc. TO1000DH, were each aligned separately to the “in silico polyploid” concatenated reference genome comprised of the B. rapa acc. R500 genome (Lou et al. 2020) with SNP correction using IMB218DH resequencing data and the B. oleracea TO1000 reference genome (Parkin et al. 2014). This was the same concatenated reference genome used to align the RNAseq data from the resynthesized lines.

We defined the expression response to homoeologous exchange as the fold change of the summed homoeolog pair expression and the parental expression, 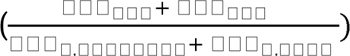. Following the approach of Coati et al. (2016) and Song et al. (2020) and calculated the coefficient of variation of this expression response 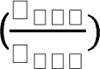 and termed it the a homoeologous exchange response variance (HERV). Statistical analysis was done with a Kruskal-Wallis test applied by the function stat_compare_means() in the R package ggpubr v.0.04.0 (R core team, 2020; Kassambara, 2020). We calculated HERV only for GO terms that contained more than 20 genes. When analyzing the response to polyploidy among different homoeolog expression biases, the expression bias of progenitor orthologs was used. The classification of biased homoeologs was taken from Bird et al. (2021) who used a cutoff of log2 fold change of 3.5 and −3.5 to classify a homoeolog as more dominantly expressed. Previous analysis showed that for over 70% of homoeologs, all six resynthesized *B. napus* lines shared the same homoeolog expression bias as the parents (Bird et al. 2021).

### Expression response to polyploidy

When investigating the dosage response to polyploidy, we limited our analysis to the syntenic homoeologous genes identified as being in a 2:2 dosage ratio. We created our dataset by combining data across individuals, selecting gene pairs in 2:2 for a particular individual sample. We did not require that a gene pair was in 2:2 dosage in every line. We calculated expression response to polyploidy for each gene pair, defined as the fold change of polyploid expression for a 2:2 syntenic homoeolog pair and the mid-parent expression of the progenitor ortholog pair 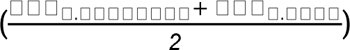. We used the same parental expression data from Bird et al. 2021 as the HERV analysis. We applied the same approach as Coate et al. (2016) and Song et al. (2020) and focused on the coefficient of variation of expression response 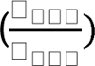 which we similarly termed the polyploid response variance (PRV). The Kruskal-Wallis implementation from ggpubr (Kassambara, 2020) was used again for statistical analysis. As for the previous analysis, we only included GO terms with 20 or more genes and defined homoeolog expression bias in terms of expression bias in parental orthologs.

## Results

### Homoeologous pairs of dosage-sensitive genes show a less variable expression response to homoeologous exchange, except when the BnA homoeolog is more highly expressed

To assess how the selection for relative dosage may affect the expression response to gene dosage changes from homoeologous exchange, we used the dosage-sensitivity gene class assignments for *Arabidopsis thaliana* from Song et al. (2020). As per Song et al. (2020), Class I Gene Ontology (GO) terms are putatively dosage-insensitive and Class II are putatively dosage-sensitive and these classes are based on the observed reciprocal retention (over-retention after whole-genome duplication, and under-retention after small-scale duplication) of genes from the investigated GO terms following the At-alpha duplication event in the Brassicaceae. Similar patterns of reciprocal retention have been identified across angiosperms. While GO terms are broad and will likely include both dosage-sensitive and -insensitive genes, Song et al. (2020) previously showed that certain GO terms result in qualitatively similar results as metabolic networks, protein-protein networks, and gene families. To leverage the superior annotation quality of *A. thaliana*, the orthologs in *B. rapa* and *B. oleracea* were assigned to the dosage-sensitivity GO classes of their *Arabidopsis* ortholog. These dosage-sensitivity assignments were used to assess how expression response differs between classes in the resynthesized allopolyploids. GO terms were then filtered so that only those with 20 or more genes in our dataset were included in the analysis

The extensive genomic rearrangements observed in this population of resynthesized lines (Xiong et al. 2011; Bird et al. 2021) provide an opportunity to test for the first time whether gene expression changes from homoeologous exchange events show signs of the constraint on relative dosage from dosage balance selection that is predicted by the Gene Balance Hypothesis. Using the published results from Bird et al. (2021), we focused on genomic regions identified as not being in 2:2 dosage, representing genomic rearrangements with 0:4, 1:3, 3:1, and 4:0 dosage ratios (BnC:BnA). To avoid the inclusion of likely aneuploidy events, genes on chromosomes that frequently showed dosage changes for the entirety or majority of the chromosome were excluded. This affected chromosome pairs 1A/1C, 2A/2C, 10A/9C (FigS1-S6). With this dataset of gene pairs affected by putative homoeologous exchange events, we compared their expression with a balanced dosage state which we represented as the summed expression of the progenitor orthologs. It should be noted, this approach did not normalize RNA with exogenous spike-in as other studies have, meaning values reported are relative gene expression levels rather than the absolute expression response. Although we can not assign expression responses to categories like dosage-dependent or compensation as previous studies have (Song et al. 2020; Shi et a. 2020; Hou et al. 2018), we can investigate the relative change in expression and test to see if it matches predictions laid out by the Gene Balance Hypothesis. This type of analysis should be robust to the issues caused by the lack of an exogenous spike-in and has been previously employed in expression comparisons of natural allopolyploid and diploid species, without the use of spike-ins (Coate et al. 2016).

We investigated the extent that expression responses from homoeologous exchanges differ among the identified dosage-sensitive and dosage-insensitive GO terms (Fig 1). We looked at the expression response of gene pairs in a given GO term, only for those gene pairs affected by a homoeologous exchange event. We used the coefficient of variation of this expression response, which we call the Homoeologous Exchange Response Variance (HERV), to assess how variable the expression response was for genes from dosage-sensitive and insensitive GO terms. After filtering GO terms with fewer than 20 genes represented in our dataset, we had 305 GO terms, with 142 dosage-insensitive and 163 dosage-sensitive. Across all lines, genes belonging to putatively dosage-sensitive GO terms showed significantly lower HERV, indicating a less variable expression response than genes from putatively dosage-insensitive GO terms (Fig 1a, Kruskal-Wallis test, p=0.00011).

**Figure 1.**
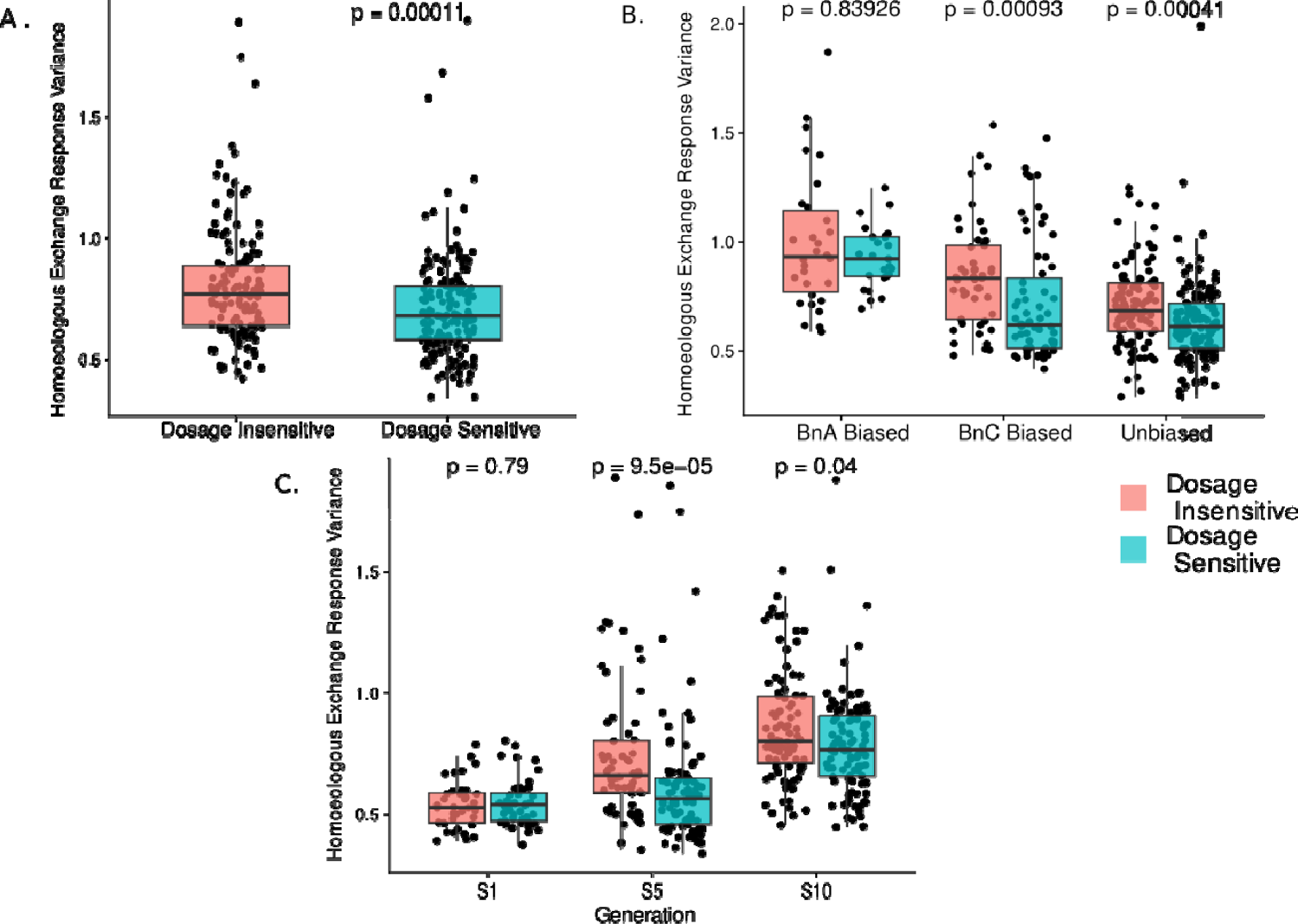
Expression changes from homoeologous exchange reflect predictions from the gene balance hypothesis Homoeologous exchange response variance (coefficient of variation of dosage response from homoeologous exchange) for all dosage imbalanced homoeologs in all 16 isogenic polyploid plants broken down by **A)** only putatively dosage-insensitive (Class I) and dosage-sensitive (Class II) GO terms from Song et al. 2020, **B)** Dosage-sensitivity classes and subgenome dominance relationship in parental lines, **C)** Dosage-sensitivity classes and generation. P-values represent results of Kruskal-Wallis test of polyploid response variance between Dosage-sensitive and insensitive GO terms. In all plots, individual dots represent a GO term, restricted only to GO terms that were represented by 20 or more genes in our dataset.

It is possible that the higher variation in expression response of genes in dosage-insensitive GO terms may be may an artifact of differences in average expression of genes in dosage-sensitive and dosage-insensitive GO terms, since there is generally lower variance for more highly expressed genes (Conesa et al., 2016; Mortazavi et al., 2008).

To rule this out we compared average TPM of homoeolog pairs for GO terms assigned as dosage-insensitive (Class I) and dosage-sensitive (Class II) with a Kruskal-Wallis test to make sure genes in dosage-sensitive GO terms did not have significantly higher expression compared to dosage-sensitive GO terms. For this dataset our results showed that dosage-sensitive (Class II) GO terms had significantly lower expression on average compared to dosage-insensitive (Class I; p=0.033; Fig S7). These results are similar to what Song et al. (2020) found for their Arabidopsis polyploids and should provide strong support that our results are not an artifact of dosage-sensitive genes being more highly expressed.

Using an allopolyploid gave us the opportunity to observe if the transcriptional response of dosage-sensitive and dosage-insensitive genes varies based on homoeolog expression bias. Such a result may suggest that the dosage constraint on homoeologous pairs from dosage balance selection can differ depending on which copy is more highly expressed. Previous transcriptomic analysis of these resynthesized lines from Bird et al. (2021) identified significantly biased homoeolog pairs defined as a log2 fold change greater than 3.5 or less than −3.5 and observed more homoeolog pairs with expression biased toward the BnC subgenome, which was dubbed the dominant subgenome (Bird et al. 2021). We compared the dosage-sensitive and dosage-insensitive GO terms, this time only including gene pairs with particular homoeolog expression bias in GO terms. This resulted in three datasets: expression response of GO terms only considering expression data for gene pairs with expression biased toward the BnC subgenome, pairs only with expression biased toward the BnA subgenome, and pairs with no expression bias. Expression bias of the gene pair was based on the expression relationship of the parental orthologs. Previous analyses by Bird et al. (2021) found these parental expression differences to match homoeolog expression bias in all six lines for over 70% of homoeologous gene pairs (Bird et al. 2021).

When broken down by direction of homoeolog expression bias there were 55 GO terms (30 dosage-insensitive and 25 dosage-sensitive), after filtering, for pairs biased toward the non-dominant BnA subgenome, 112 GO terms (49 dosage-insensitive and 63 dosage-sensitive) for pairs biased toward the dominant BnC subgenome, and 239 (105 dosage-insensitive and 134 dosage-insensitive) for gene pairs without expression bias. We found that homoeologous gene pairs with expression biased toward the dominant BnC subgenome (Kruskal-Wallis test, p=0.00093) and unbiased gene pairs (Kruskal-Wallis test, p=0.00041) show significantly lower HERV in dosage-sensitive GO terms than dosage-insensitive GO terms while pairs with expression biased toward the BnA subgenome did not show a significant difference between classes (Fig 1b; Kruskal-Wallis test, p=0.83926). Thus, we present evidence that the expression response of dosage-sensitive gene pairs differs depending on which homoeolog is more highly expressed.

When analyzing expression response by generation, there were 80 GO terms (36 dosage-insensitive and 44 dosage-sensitive) that passed filtering for generation one, 148 (63 dosage-insensitive and 85 dosage-sensitive) for generation five, and 187 (87 dosage-insensitive and 100 dosage-insensitive) for generation 10. We found that there was not a significant difference in HERV between dosage-sensitive and dosage-insensitive GO terms at the first generation (Fig 1c, Kruskal-Wallis test, p=0.79), but dosage-sensitive and insensitive GO terms did show different HERV at the fifth and tenth (Fig 2c, Kruskal-Wallis test, p=9.5×10^-5^, p=0.04). We also found that homoeologous exchange response variance increased over time with dosage-sensitive and dosage-insensitive GO terms showing mean HERV of 0.661 and 0.502, respectively, in generation one and increasing to 0.828 and 0.697, respectively, in generation ten.

**Fig 2.**
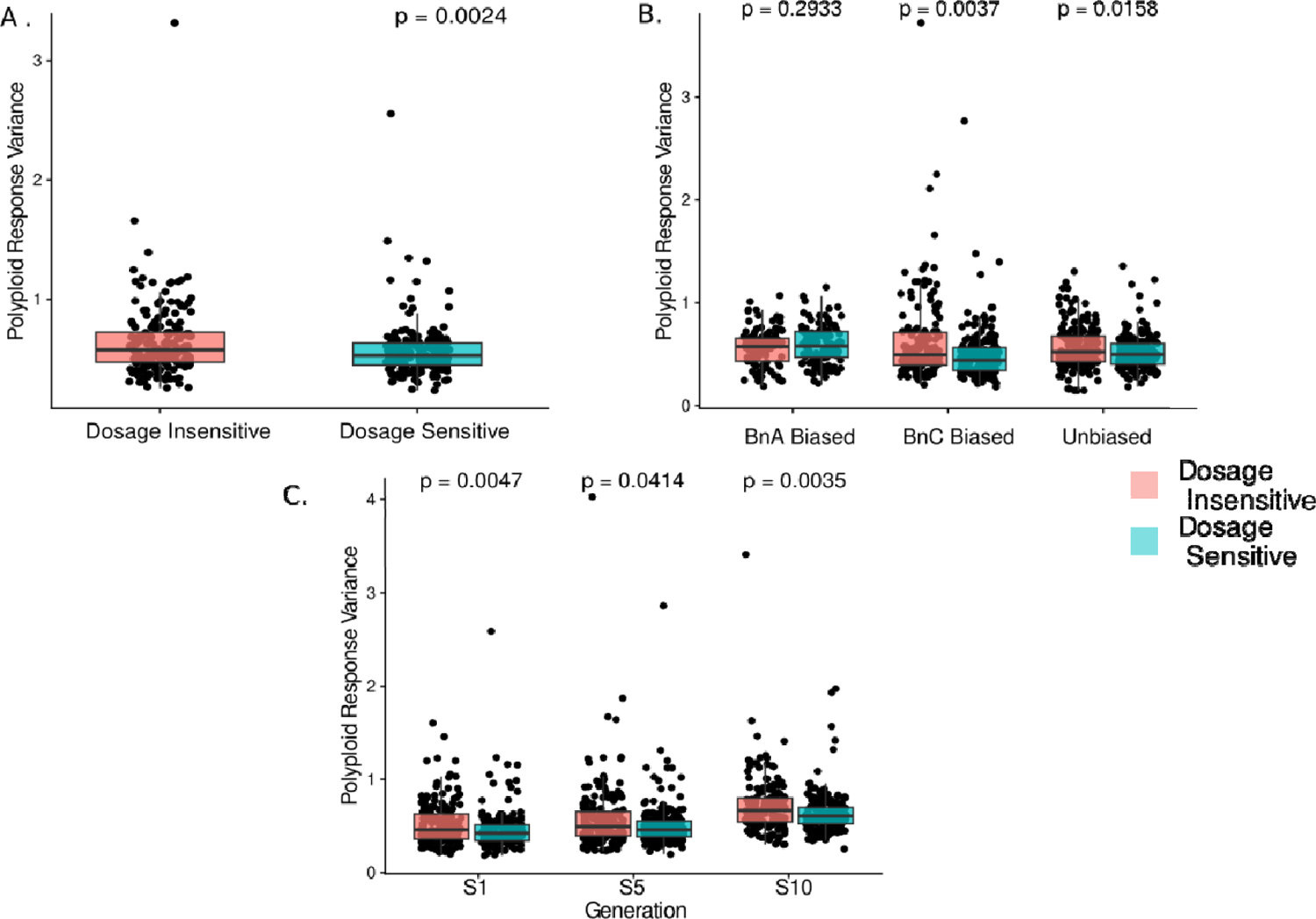
Expression changes from allopolyploidy reflect predictions from the gene balance hypothesis Polyploid response variance (coefficient of variation of dosage response) for all 2:2 balanced homoeologs in all 16 isogenic polyploid plants broken by **A)** only putatively dosage-insensitive (Class I) and dosage-sensitive (Class II) GO terms from Song et al. 2020, **B)** Dosage-sensitivity classes and subgenome dominance relationship in parental lines, **C)** Dosage-sensitivity classes and generation. P-values represent results of Kruskal-Wallis test of polyploid response variance between Dosage-sensitive and insensitive GO terms. In all plots, individual dots represent a GO term, restricted only to GO terms that were represented by 20 or more genes in our dataset.

### Homoeologous pairs of dosage-sensitive genes show less variable expression response to WGD, except when the BnA homoeolog is more highly expressed

We further investigated the transcriptional response to dosage changes and variation by subgenome expression bias by analyzing the relative gene expression change for individual homoeologous gene pairs in 2:2 dosage. We took the fold change of the summed transcript count for homoeologous gene pairs in the allopolyploid individuals and mid-parent value of the progenitor orthologs. We used the polyploid response variance (PRV) measure from Song et al. (2020) and Coate et al. (2016), defined as the coefficient of variation of the relative expression response, to assess how variable the expression response to polyploidy is in the different gene groups. These analyses allowed us to establish the expression response to polyploidy in a newly formed allopolyploid and further explore and validate the findings about homoeolog expression bias in the HERV analysis.

Analyzing data across all lines and filtering out GO terms with fewer than 20 genes, we had a final count of 376 GO terms of which 181 were classified dosage-insensitive and 195 were dosage-sensitive. As observed previously in resynthesized autopolyploids and natural *Glycine* allopolyploids, the polyploid response variance was significantly lower (i.e. the expression response was less variable) in genes from GO terms in the dosage-sensitive class compared to the dosage-insensitive class (Kruskal-Wallis test, p=0.0024; Fig 2; Fig 2a). We again checked for expression differences between dosage-sensitive and dosage-insensitive groups of genes. For this dataset of homoeologous pairs at 2:2 dosage, our results showed that dosage-sensitive (Class II) GO terms had significantly lower expression on average compared to dosage insensitive (Class I; p=0.0085; Fig S8). This again supports that our results are not due to differences in expression between genes in the Class I and Class II GO terms.

We next sought to replicate the above results showing a less variable expression response for dosage-sensitive gene pairs when the BnA homoeolog was more highly expressed, this time using PRV. After filtering out GO terms with fewer than 20 genes, there were 274 GO terms (113 dosage-insensitive, 124 dosage-sensitive) for gene pairs biased toward the non-dominant BnA subgenome, 330 GO terms (156 dosage-insensitive and 174 dosage-sensitive) for pairs biased toward the dominant BnC subgenome, and 374 GO terms (179 dosage-insensitive and 195 dosage-sensitive) for genes not biased toward either subgenome. We again found that pairs with expression biased toward the *B. napus* C subgenome (BnC), or with unbiased expression, showed a significant difference between PRV of dosage-sensitive and dosage-insensitive GO terms as above (Kruskal-Wallis test, p=0.0037; p=0.0158, respectively). As before, gene pairs biased toward the *B. napus* A subgenome (BnA) showed no significant difference in PRV between dosage-sensitive and insensitive GO classes (Kruskal-Wallis test, p=0.2933; Fig 2b). These results provide further support that the expression response of dosage-sensitive genes differs depending on which homoeolog is more highly expressed.

When broken down by generation, there were 375 GO terms (180 dosage-insensitive and 195 dosage-sensitive) that passed filtering for generation one, 368 GO terms (174 dosage-insensitive and 194 dosage-sensitive) for generation five, and 362 GO terms (172 dosage-insensitive and 190 dosage-sensitive) for generation ten. In all three generations, dosage-sensitive GO terms have significantly lower PRV than dosage-insensitive GO terms (Fig 2c; Gen 1 p=0.0047, Gen 5 p=0.0414, Gen 10 p=0.0035). We observed an increase in the coefficient of variation over time, with both dosage-sensitive and dosage-insensitive showing higher PRV in generation ten than in the first generation (Fig 2c). Notably, in generation ten the dosage-sensitive GO terms show higher mean polyploidy response variance than dosage-insensitive GO terms in the first generation.

### Expression response to homoeologous exchange is more variable than the expression response to WGD

The Gene Balance Hypothesis predicts that dosage changes that alter only a subset of genes in a pathway or protein complex will produce genomic imbalance that can reduce organismal fitness, while changes that alter the dosage of the entire genome will maintain the necessary stoichiometric balance of gene products. Given that HEs result in dosage and expression changes for only some regions of the genome, they are predicted to produce genomic imbalance. While we do not have phenotypic data to directly investigate genomic imbalance, previous work has shown a greater variation in expression response to aneuploidy than polyploidy in *Arabidopsis thaliana* (Hou et al. 2018). These authors argued the patterns of expression modulation exhibited by aneuploids reflected the same principles of genomic balance as is observed at the phenotypic level. Therefore, investigating patterns of expression between genes affected by HE and polyploidy may still shed light on whether HEs produce genomic imbalance. To test for signs of genomic imbalance, we compared the coefficient of variation for the expression response to the polyploidy and homoeologous exchange datasets to see if the HE expression response was more variable (Fig 3).

**Figure 3.**
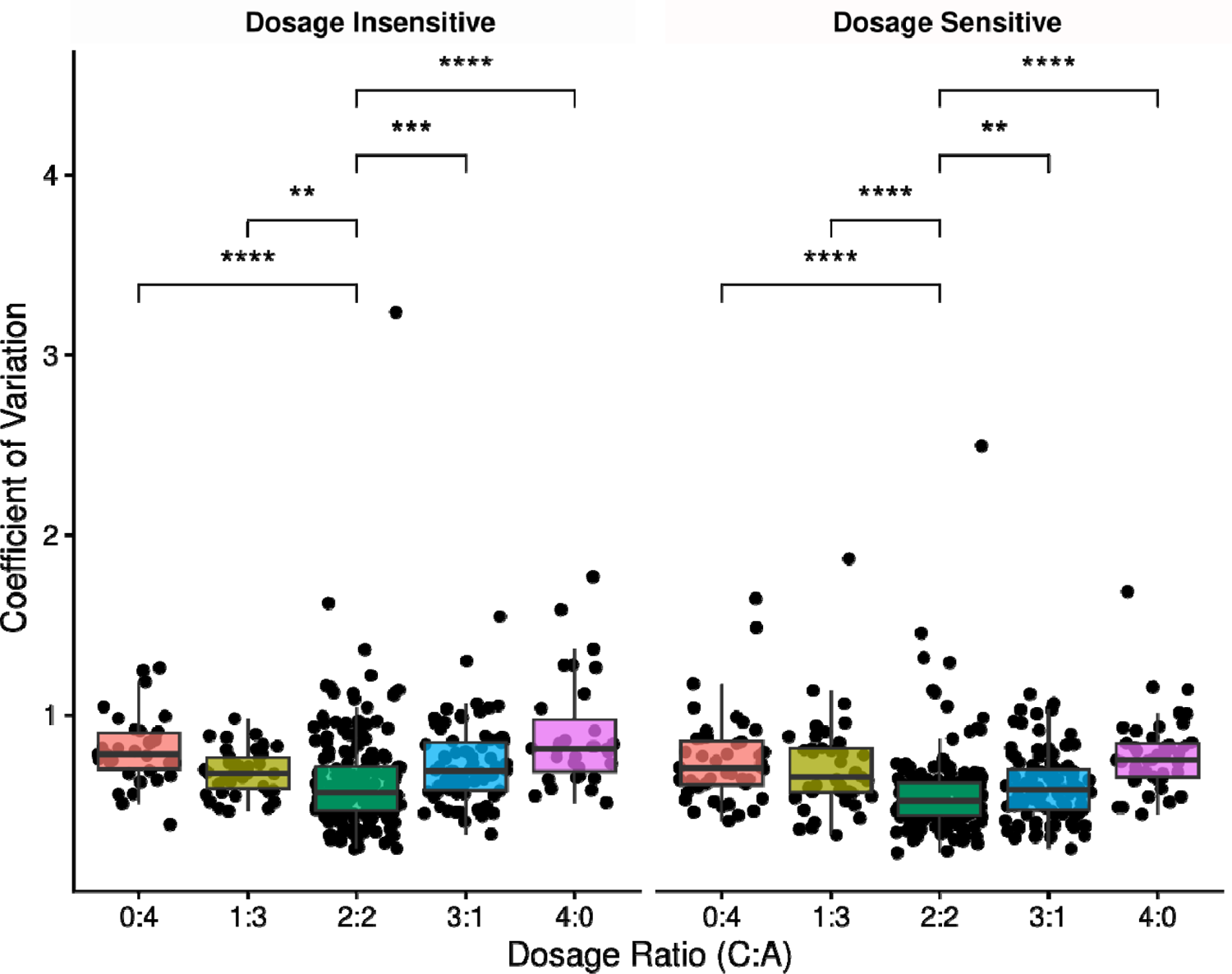
Expression responses to homoeologous exchange is more variable than the response to polyploidy Comparison of the coefficient of variation for the expression response to homoeologous exchanges (dosages 0:4, 1:3, 3:1, and 4:0) and allopolyploidy (dosage 2:2) for dosage insensitive GO terms (left) and dosage-sensitive GO terms (right) combined across all lines and generations. Individual dots represent a GO term, restricted only to GO terms that were represented by 20 or more genes in our dataset. Asterisks represent p-value of pairwise Wilcoxon tests between an HE dosage group (0:4, 1:3, 3:1, 4:0) and the 2:2 group, representing the polyploidy expression response. Asterisks represent significance levels as follows: “*” = p ≤ 0.05, “**” = p ≤ 0.01, “***” = p ≤ 0.001, “****” = p ≤ 0.0001

First, we compared the proportion of gene pairs belonging to dosage-sensitive and dosage-insensitive GO terms in all 16 individuals for the polyploidy and homoeologous exchange analysis. For the polyploid analysis, the mean proportion of genes belonging to dosage-insensitive GO terms is 0.554, while it is 0.544 for the homoeologous exchange analysis. This difference in the proportion of Class I and Class II GO terms between the PRV and HERV analysis was not statistically significant (l-^2^-test, p=0.983). Even considering the direction of difference in the proportion of dosage-insensitive genes, a greater proportion of gene pairs having dosage-insensitive GO terms would be predicted to result in a higher coefficient of variation. Instead, we found a significantly higher coefficient of variation from all pairwise comparisons between the expression response to homoeologous exchanges of different directions and dosage and the expression response to polyploidy. This was the case for both dosage-sensitive and dosage-insensitive GO terms. (Fig 3, Wilcoxon test, p < 0.01). Additionally, the results appear to show an additive pattern where the coefficient of variation in expression response becomes larger and more significantly different from the coefficient of variation of the polyploidy expression response for the 0:4 and 4:0 HEs, which represent larger dosage changes than 1:3 and 3:1 HEs. These results further support the prediction that HEs can create genomic imbalance, because a more variable expression response is predicted to translate to more variation in the final amount of interacting gene products, which disrupts the stoichiometry of the system.

Finally, the significantly different expression response to homoeologous exchanges and polyploidy-induced dosage changes in all comparisons is strong evidence that the patterns observed for homoeologous exchange-induced dosage changes are, at least partially, distinct from the effects of polyploidy-induced dosage change and our results from previous sections are not merely an artifact of our HE analysis picking up the effects of dosage changes caused by allopolyploidy or trans dosage effects of aneuploidy.

## Discussion

### Genomic balance and the evolution of homoeologous exchanges

Homoeologous exchanges have long been recognized as an engine of phenotypic diversity and novelty in newly formed polyploids (Pires et al. 2004; Gaeta et al. 2007; Xiong et al. 2011; Rousseau-Guetin et al. 2017; Stein et al. 2017; Leal-Bertioli et al. 2018; Lloyd et al. 2018; Mason and Wendel 2020; Ferreira de Carvalho et al 2021, Wu et al. 2021; Bomblies 2023). Our analysis of genomic rearrangements and homoeologous exchanges in resynthesized *B. napus* confirmed at higher resolution the extensive rearrangements in these lines (Gaeta et al. 2007; Xiong et al. 2011). Investigations of genomic balance and dosage-sensitivity have predominantly focused on polyploidy and aneuploidy as the sources of gene dosage alteration (Hou et al. 2018; Yang et al. 2021; Shi et al. 2021). However, homoeologous exchanges, which alter the dosage ratio of parental chromosome segments, have also been shown to produce dosage-dependent expression changes that perturb the total expression of a gene pair (Lloyd et al. 2018). We tested two predictions from the Gene Balance Hypothesis regarding the transcriptional response to homoeologous exchanges. The first is that the dosage balance selection to maintain the stoichiometry of gene products will result in dosage-sensitive genes having a less variable expression response to homoeologous exchange. The second is that, due to the perturbation to stoichiometry from homoeologous exchanges, the expression response to homoeologous exchanges will be more variable than the expression response to whole-genome duplication.

We found that expression response to homoeologous exchanges matches those previously observed in response to whole-genome duplication (Fig 1a) (Coate et al. 2016; Song et al. 2020). Gene expression responses from dosage-sensitive GO terms are less variable than those from dosage-insensitive GO terms, as predicted by the Gene Balance Hypothesis. However, we also saw the difference in expression response between dosage-sensitive and dosage-insensitive genes was not present for homoeolog pairs with expression biased toward the non-dominant BnA subgenome. When comparing expression response variation from homoeologous exchanges to polyploidy, we observe significantly higher variation in the expression response to HEs (Fig 3). Similar results from comparisons of expression modulation from aneuploidy and polyploidy have been taken as evidence for genomic imbalance, and an ultimate explanation for the cause of the greater phenotypic impacts and fitness cost of aneuploidy (Hou et al. 2018; Shi et al. 2020). These results similarly support a genomic imbalance arising from HEs and raise the possibility that a similar fitness cost due to genomic imbalance exists for HEs, though the magnitude relative to other fitness costs like meiotic instability cannot be determined here. Such results have not been reported before, to our knowledge, and provide strong evidence that expression response of dosage-sensitive genes to homoeologous exchanges is affected by dosage balance selection.

If homoeologous exchanges evolve in ways predicted by the Gene Balance Hypothesis then we might expect dosage balance selection to disfavor homoeologous exchanges containing dosage-sensitive genes, producing biases in the gene functions surviving homoeologous exchanges that are similar to those for small-scale duplications. Indeed, Hurgobin et al. (2018) and Bayer et al. (2021) identified a significant degree of gene presence-absence variation in *B. napus* arising from homoeologous exchanges. Presence-absence variation was negatively associated with membership in protein-protein interaction networks (Bayer et al. 2021) and positively associated with GO terms related to plant defense and stress pathways (Hurgobin et al. 2018). They also observed several homoeologous exchanges generating presence-absence variation in paralogs of the large gene family *FLC,* which regulates flowering time. Analysis of expression dynamics of *FLC* paralogs in *B. napus* showed that while FLC paralogs are dosage-sensitive, the selection on dosage balance acts on overall *FLC* gene family expression allowing compensatory drift (Thompson et al. 2016) and expression divergence (Calderwood et al. 2020). This *FLC* example shows that the interplay of homoeologous exchange and dosage balance may be highly dynamic depending on the gene family in question. The effect of dosage-sensitivity on expression response to HEs observed here may also help understand the mechanisms reported by Lloyd et al. (2018), which observed a tendency of transcriptional compensation of older HE events in natural *B. napus*.

Finally, selection for dosage balance may also drive subgenome biases in the direction of homoeologous exchange. For example, Edger et al. (2019) proposed that the selection for stoichiometric balance of gene products could explain the overwhelming bias in direction of homoeologous exchange, favoring the dominant subgenome, in the octoploid strawberry genome. These results might help explain why natural *B. napus* lines tend to show bias in favor of HE events decreasing copies of C subgenome regions (Nicolas et al. 2012; Chalhoub et al. 2014), as perturbing the level of gene products among only a fraction of the members of interacting units is expected to incur a fitness cost similar to aneuploidy. Our results show that the effect of dosage-sensitivity on the transcriptional response to dosage changes differs for gene pairs depending on which homoeolog is more highly expressed. If the less variable expression response of dosage-sensitive genes is truly a result of the dosage balance selection on homoeologous pairs to maintain proper stoichiometry of their gene products this raises the possibility that dosage balance selection acts on homoeologous pairs differently depending on which copy is more highly expressed. This pattern suggests that the changes to homoeologs from one subgenome, in this case the dominant BnC subgenome, are more likely to contribute to stoichiometric imbalance.

It is worth noting, we were not able to completely rule out expression changes from other gene loss or silencing processes, like partial chromosomal deletion or duplication or DNA methylation. However, it is unlikely that our results are driven primarily by these other factors. Gaeta et al. 2007 analyzed genomic, epigenomic and transcriptomic changes in this population in the 5th generation using gel-based markers, where 71% of genetic RFLP marker deletions were accompanied by intensifications by homoeologous markers from the other subgenome, supporting homoeologous exchange events in the majority of cases. Gaeta et al. 2007 also reported no correlation between epigenomic markers and expression changes, measured by cDNA-AFLP markers, in the siblings of this population. Similarly, Hou et al. (2018) looked at gene expression and DNA methylation changes in an aneuploid series of Arabidopsis and did not identify consistent changes in DNA methylation in any context from aneuploidy in any context that suggested genomic imbalance and its expression effects were mechanistically caused by methylation changes. Still, future work combining cytogenetics, genomics, and epigenomics more directly will provide a clearer picture of the kinds of expression patterns reported here and other factors like non-HE genomic rearrangement and epigenetic changes.

### The effect of dosage-sensitivity on expression responses differs depending on which subgenome is more highly expressed

In both our HERV and PRV analyses, we observed evidence of the difference in expression respothe nse between dosage-sensitive and dosage-insensitive genes being absent for homoeologous pairs with expression biased toward the non-dominant subgenome. This may indicate that the constraint from dosage balance selection differs between dosage-sensitive gene pairs depending on the direction of their homoeolog expression bias. Previous analysis of these resynthesized lines showed that homoeologous pairs biased toward the BnC subgenome, which was the maternal contributor and called the dominant subgenome, were more connected in a protein-protein interaction network, while pairs with expression biased toward the paternal BnA subgenome showed no such enrichment for connectivity (Bird et al. 2021). This lack of connectivity may explain why putatively dosage-sensitive genes with biased expression toward the non-dominant subgenome do not show less variable expression; without high connectivity in gene networks, they do not experience strong dosage-balance selection. Bird et al. (2021) speculated that this enrichment was driven by interactions between the nuclear and organellar genomes, given the functional enrichment for mitochondria, chloroplast, and cytoplasm among the PPI network, however, some recent work casts doubt on the impact of cytonuclear incompatibilities on this kind of response to allopolyploidy (Ferreira de Carvalho et al. 2019; Sharborough et al. 2022). Assessing the generality of these subgenome differences in network connectivity and their relation to cytonuclear interaction will be a promising avenue for future research in this area.

It is noteworthy that we see no difference in the expression response of dosage-sensitive genes and dosage-insensitive genes for both HEs and whole-genome duplication. This suggests it is a general aspect of how homoeolog expression bias and dosage-sensitive affect the expression response to dosage changes. Given the hypothesized importance of coordinated expression responses for maintaining the stoichiometry of gene products after dosage changes (Coate et al. 2016; Song et al. 2020), this may have implications for differences in long-term duplicate gene retention patterns between subgenomes. For example, over the long term, subgenome differences in expression responses might be predicted to preserve more dosage-sensitive genes from the dominant subgenome than the non-dominant since dosage-sensitive gene pairs with expression biased toward the dominant subgenome will be predicted to be retained more. In line with this, Schnable et al. (2012) observed that biased retention of dosage-sensitive genes broke down over time, with only 50% of genes retained from one genome duplication event being retained in duplicate after a subsequent duplication event. They further observed that the lower expressed copy was more likely to be lost and proposed the lower expressed copies contribute less to final interacting gene products, and so experience less purifying selection and weaker dosage constraint (Schnable et al. 2012). Similarly, when subgenome dominance was first described in *Arabidopsis*, the dominant subgenome was also associated with clusters of dosage-sensitive genes across the genome (Thomas et al. 2006).

### Implications for long-term duplicate gene evolution and the interplay of biased fractionation and reciprocal retention of duplicate genes

We propose a unified model for short-term and long-term interactions of subgenome dominance and dosage balance. This model involves: (1) greater retention of dosage-sensitive gene pairs that are biased toward the dominant subgenome due to stronger dosage balance selection and (2) the eventual divergence of duplicates over long evolutionary time and loss of non-dominant homoeologs due to biased fractionation. Following duplication, gene pairs biased toward the dominant subgenome in these synthetic *B. napus* show higher connectivity in protein-protein interaction networks and functional enrichment. Dosage-sensitivity is a spectrum, most strongly correlated with the connectivity of gene products in a network or macromolecular complex (Veitia and Birchler, 2012). Therefore, the less variable expression response of unbiased and dominant subgenome-biased gene pairs may be reflective of greater dosage sensitivity than pairs biased toward the non-dominant subgenome. Greater dosage sensitivity predicts that these gene pairs will be retained for a longer time due to dosage balance selection to maintain proper stoichiometry, given that their loss is more likely to perturb the relative balance of interacting gene products (Freeling et al. 2012).

Dosage constraints on gene duplicates are not permanent and can change or subside over evolutionary time (Bekaert et al. 2011; Schnable et al. 2012; Conant et al. 2014). Additionally, the stoichiometry of interacting proteins is what is truly under dosage balance selection. The expression of individual paralogs can diverge so long as that stoichiometry is largely left intact, a phenomenon called compensatory drift (Thompson et al. 2016). In the case of subgenome dominance, one copy is contributing a greater fraction of the overall amount of that gene product. As dosage constraint weakens, deleting the more highly expressed copy will cause a greater disturbance to the stoichiometric balance (Freeling et al. 2012). As an extreme example, if one copy contributes 90% to total expression and the other 10%, a greater stoichiometric imbalance would be observed with interactors when losing the dominant (90%) copy. As such, the dominant copy will be under stronger purifying selection. Under compensatory drift it is easier for the dominant copy to change expression enough to account for all or most of the gene product of a pair, thus reducing purifying selection on the non-dominant copy which now contributes little-to-none to the stoichiometry balance.

This difference in purifying selection reduces the likelihood that the dominant copy is fractionated by the short-deletion mechanism postulated to drive genome fractionation in plant genomes (Woodhouse et al. 2010). Ultimately, genes on the non-dominant subgenome will be preferentially lost and the dominant subgenome will maintain higher gene content and more enrichment for dosage-sensitive genes - even through successive polyploid events (Woodhouse et al., 2014).

### Future Directions

Several findings may warrant follow-up or more targeted investigation. Our comparison of homoeologous exchange and polyploidy response variance showed that overall gene expression was more variable in response to homoeologous exchange compared to polyploidy. The concept that selection for beneficial HE’s may be limited by a polyploid ratchet (Gaeta and Pires 2010) or other constraints related to the negative effect of HEs on meiotic stability has been discussed in the context of allopolyploid systems such (e.g., Leal-Bertioli et al. 2018, Lloyd et al. 2018, Pele et al. 2018, Edger et al. 2019, Ferreira de Carvalho et al. 2021, Higgins et al. 2021, Wu et al. 2021, Deb et al. 2023). Our results expand on these findings by suggesting HEs may also be selected against due to the genomic imbalance they produce. Replicating these results beyond gene classes based on GO terms, such as more fine-resolution data like metabolic pathway membership, protein-protein interaction networks, or gene families may provide more precise estimates of effects, as functional classification based on GO terms is known to introduce heterogeneity. Additionally, this study only looks at expression in leaf tissue. Expression differences and differences in homoeolog expression bias are likely to exist across tissues. Investigation in more tissue types and looking for variation by tissue is a promising avenue of research. Future work would also benefit from approaches using spike-ins and those which can isolate homoeologous exchange from other trans-effects on expression, both from hybridization and aneuploidy, that could not be controlled for when assessing expression changes in this study. Such work would provide more precise estimates of the magnitude of the effect of dosage-sensitivity on expression responses to dosage changes. This improved experimental design would also help make sense of the finding that PRV and HERV for both dosage-sensitive and -insensitive GO terms increase over time (Fig 1c, Fig 2c). Currently, it is not possible to distinguish whether this result is from changes in the strength of dosage constraints or an accumulation of inter-individual variation from trans-dosage effects in the genomic background. Disentangling these two explanations will reveal novel insights into the dynamics of expression changes and genomic balance over short evolution time scales. One particularly interesting possibility would be exploring ways to generate or introduce homoeologous exchanges of a specific dosage in a controlled genetic background, allowing a more precise investigation of the effect of dosage changes and the transcriptional response.

## Conclusion

This study provides new evidence on the potential for genomic imbalance from homoeologous exchanges and insight into how dosage balance selection affects the gene expression changes from genomic rearrangements. These findings may help fuel more integrative genetic and evolution investigations of homoeologous exchange, subgenome expression dominance, and duplicate gene evolution that can leverage the vast new output of genomes with ancestral and recent polyploidy and explicit evolutionary models of ancestral subenomes (Emery et al. 2018; Hao et al. 2021, 2022; Parey et al. 2022). This new avenue of investigation may help further examine evolution and epistasis as well as selection and divergence among paralogs (Qi et al. 2021; Conover and Wendel, 2022; Kwon et al. 2022), and spur further integration of methods and data across phylogenomics, comparative and population genomics, and network biology (Renny-Byfield et al. 2017; Blischak et al. 2018). Such work can enhance plant breeding efforts by providing a strengthened evolutionary understanding of the consequences of gene duplicates, structural variation, relative gene dosage, and subgenome dominance (Rodriguez-Leal et al. 2017; Bird et al. 2018; Salman-Minkov et al. 2016; Turner-Hissong et al. 2020; Bayer et al. 2021; Bomblies 2023).

## Data availability

Raw data from this project are available on the NCBI Sequence Read Archive (SRA) Project PRJNA577908. Intermediate files can be found at https://doi.org/10.5061/dryad.h18931zjr and code to recreate main figures and results can be found at https://github.com/KevinABird/Bird_GenomeInFlux_BNapus

## Acknowledgments

The authors would like to thank the editors and 3 anonymous reviewers for helpful and constructive comments on earlier versions of this manuscript. This work was supported by MSU-AgBioResearch funding to PPE and RV, USDA-NIFA HATCH 1009804 and NSF-IOS PGRP 2029959 to PPE, NSF-GRFP DGE-1424871, NSF-IOS PRFB 2208944 to KAB and National Key Research and Developmental Program of China (2016YFD0100202) and National Natural Science Foundation of China (31871239 and 31471173) to ZX.

## Author Contributions

KAB contributed to conceptualization, formal analysis, investigation, methodology, validation, visualization, writing—original draft, and writing—review editing; JCP. contributed to resources, and writing—review editing; RV. contributed to supervision, methodology, and writing—review editing; ZX. contributed resources, experimental design, and writing—review editing; and PPE. contributed supervision, methodology, and writing – review editing.

## References

Alger, E. I., & Edger, P. P. (2020). One subgenome to rule them all: underlying mechanisms of subgenome dominance. Current Opinion in Plant Biology, 54, 108–113.

Bekaert, M., Edger, P. P., Pires, J. C., & Conant, G. C. (2011). Two-phase resolution of polyploidy in the *Arabidopsis* metabolic network gives rise to relative and absolute dosage constraints. The Plant Cell, 23(5), 1719–1728.

Bayer, P. E., Scheben, A., Golicz, A. A., Yuan, Y., Faure, S., Lee, H., … & Edwards, D. (2021). Modelling of gene loss propensity in the pangenomes of three *Brassica* species suggests different mechanisms between polyploids and diploids. Plant Biotechnology Journal, 19(12), 2488–2500.

Birchler, J. A., Bhadra, U., Bhadra, M. P., & Auger, D. L. (2001). Dosage-dependent gene regulation in multicellular eukaryotes: implications for dosage compensation, aneuploid syndromes, and quantitative traits. Developmental Biology, 234(2), 275–288.

Birchler, J. A., & Newton, K. J. (1981). Modulation of protein levels in chromosomal dosage series of maize: the biochemical basis of aneuploid syndromes. Genetics, 99(2), 247–266.

Birchler, J. A., & Veitia, R. A. (2007). The gene balance hypothesis: from classical genetics to modern genomics. The Plant Cell, 19(2), 395–402.

Birchler, J. A., & Veitia, R. A. (2010). The gene balance hypothesis: implications for gene regulation, quantitative traits and evolution. New Phytologist, 186(1), 54–62.

Birchler, J. A., & Veitia, R. A. (2012). Gene balance hypothesis: connecting issues of dosage sensitivity across biological disciplines. Proceedings of the National Academy of Sciences, 109(37), 14746–14753.

Bird, K. A., Niederhuth, C. E., Ou, S., Gehan, M., Pires, J. C., Xiong, Z., … & Edger, P. P. (2021). Replaying the evolutionary tape to investigate subgenome dominance in allopolyploid *Brassica napus*. New Phytologist, 230(1), 354–371.

Bird, K. A., VanBuren, R., Puzey, J. R., & Edger, P. P. (2018). The causes and consequences of subgenome dominance in hybrids and recent polyploids. New Phytologist, 220(1), 87–93.

Blanc, G., & Wolfe, K. H. (2004). Functional divergence of duplicated genes formed by polyploidy during *Arabidopsis* evolution. The Plant Cell, 16(7), 1679–1691.

Blischak, P.D, Mabry, M.E., Conant, G.C., and Pires, J.C. (2018). Integrating networks, phylogenomics, and population genomics for the study of polyploidy. Annual Review of Ecology, Evolution, and Systematics 49: 253–278.

Blischak, P.D., Sajan, M., Barker, M.S., & Gutenkunst, R.N. (2023). Demographic history inference and the polyploid continuum. bioRxiv, https://doi.org/10.1101/2022.09.15.508148

Bomblies, K. (2023). Learning to tango with four (or more): the molecular basis of adaptation to polyploid meiosis. Plant Reproduction, 36(1), 107–124.

Cao, Y., Zhao, K., Xu, J., Wu, L., Hao, F., Sun, M., … & Xiong, Z. (2023). Genome balance and dosage effect drive allopolyploid formation in *Brassica*. Proceedings of the National Academy of Sciences, 120(14), e2217672120.

Calderwood, A., Lloyd, A., Hepworth, J., Tudor, E. H., Jones, D. M., Woodhouse, S., … & Morris, R. J. (2021). Total FLC transcript dynamics from divergent paralogue expression explains flowering diversity in *Brassica napus*. New Phytologist, 229(6), 3534–3548.

Chalhoub, B., Denoeud, F., Liu, S., Parkin, I. A., Tang, H., Wang, X., … & Wincker, P. (2014). Early allopolyploid evolution in the post-Neolithic *Brassica napus* oilseed genome. Science, 345(6199), 950–953.

Chawla, H.S., Lee, H.T., Gabur, I., Vollrath, P. Tamilselvan-nattar-Amutha, S, … & Snowdon, R.J. (2021). Long-read sequencing reveals widespread intragenic structural variants in a recent allopolyploid crop plant. Plant Biotechnology Journal 19(2), 240–250

Chen, Z. J. (2007). Genetic and epigenetic mechanisms for gene expression and phenotypic variation in plant polyploids. Annual Review of Plant Biology, 58, 377–406.

Cheng, F., Wu, J., Fang, L., Sun, S., Liu, B., Lin, K., … & Wang, X. (2012). Biased gene fractionation and dominant gene expression among the subgenomes of *Brassica rapa*. PloS One, 7(5), e36442.

Cheng, F., Sun, C., Wu, J., Schnable, J., Woodhouse, M.R., Liang, J., Cai, C., Freeling, M. and Wang, X. (2016), Epigenetic regulation of subgenome dominance following whole genome triplication in *Brassica rapa*. New Phytologist, 211 (1), 288–299. https://doi.org/10.1111/nph.13884

Coate, J. E., Song, M. J., Bombarely, A., & Doyle, J. J. (2016). ExpressionlJlevel support for gene dosage sensitivity in three *Glycine* subgenus *Glycine* polyploids and their diploid progenitors. New Phytologist, 212(4), 1083–1093.

Conant, G. C., Birchler, J. A., & Pires, J. C. (2014). Dosage, duplication, and diploidization: clarifying the interplay of multiple models for duplicate gene evolution over time. Current Opinion in Plant Biology, 19, 91–98.

Conesa, A., Madrigal, P., Tarazona, S., Gomez-Cabrero, D., Cervera, A., McPherson, A., … & Mortazavi, A. (2016). A survey of best practices for RNA-seq data analysis. Genome biology, 17(1), 1–19.

Conover, J.L., & Wendel, J.F. (2022). Deleterious mutations accumulate faster in allopolyploid than diploid cotton (*Gossypium*) and unequally between subgenomes. Molecular Biology and Evolution, 39(2) msac024

De Smet, R., Adams, K. L., Vandepoele, K., Van Montagu, M. C., Maere, S., & Van de Peer, Y. (2013). Convergent gene loss following gene and genome duplications creates single-copy families in flowering plants. Proceedings of the National Academy of Sciences, 110(8), 2898–2903.

Deb, S. K., Edger, P. P., Pires, J. C., & McKain, M. R. (2023). Patterns, mechanisms, and consequences of homoeologous exchange in allopolyploid angiosperms: A genomic and epigenomic perspective. New Phytologist. 238: 2284–2304. https://doi.org/10.1111/nph.18927

Edger, P. P., & Pires, J. C. (2009). Gene and genome duplications: the impact of dosage-sensitivity on the fate of nuclear genes. Chromosome Research, 17(5), 699–717.

Edger, P. P., Poorten, T. J., VanBuren, R., Hardigan, M. A., Colle, M., McKain, M. R., … & Knapp, S. J. (2019). Origin and evolution of the octoploid strawberry genome. Nature Genetics, 51(3), 541–547.

Edger, P. P., Smith, R., McKain, M. R., Cooley, A. M., Vallejo-Marin, M., Yuan, Y., … & Puzey, J. R. (2017). Subgenome dominance in an interspecific hybrid, synthetic allopolyploid, and a 140-year-old naturally established neo-allopolyploid monkeyflower. The Plant Cell, 29(9), 2150–2167.

Emery, M., Willis, M.M.S., Hao, Y., Barry, K., Oakgrove, K., Peng, Y., Schmutz, J., Lyons, E., Pires, J.C., Edger, P.P., and Conant, G.C. (2018). Preferential retention of genes from one parental genome after polyploidy illustrates the nature and scope of the genomic conflicts induced by hybridization. PLOS Genetics 14(3): e1007267.

Ferreira de Carvalho, J., Lucas, J., Deniot, G., Falentin, C., Filangi, O., Gilet, M., Legeai, F., Lode, M., Morice, J., Trotoux, G., Aury, J.-M., Barbe, V., Keller, J., Snowdon, R., He, Z., Denoeud, F., Wincker, P., Bancroft, I., Chèvre, A.-M. and Rousseau-Gueutin, M. (2019), Cytonuclear interactions remain stable during allopolyploid evolution despite repeated whole-genome duplications in *Brassica*. Plant Journal, 98(3), 434–447.

Ferreira de Carvalho, J., Stoeckel, S., Eber, F., LodélJTaburel, M., Gilet, M. M., Trotoux, G., … & Rousseau-Gueutin, M. (2021). Untangling structural factors driving genome stabilization in nascent *Brassica napus* allopolyploids. New Phytologist, 230(5), 2072–2084.

Freeling, M. (2009). Bias in plant gene content following different sorts of duplication: tandem, whole-genome, segmental, or by transposition. Annual Review of Plant Biology, 60, 433–453.

Freeling, M., & Thomas, B. C. (2006). Gene-balanced duplications, like tetraploidy, provide predictable drive to increase morphological complexity. Genome Research, 16(7), 805–814.

Freeling, M., Woodhouse, M. R., Subramaniam, S., Turco, G., Lisch, D., & Schnable, J. C. (2012). Fractionation mutagenesis and similar consequences of mechanisms removing dispensable or less-expressed DNA in plants. Current Opinion in Plant Biology, 15(2), 131–139.

Gaeta, R. T., Pires, J. C., Iniguez-Luy, F., Leon, E., & Osborn, T. C. (2007). Genomic changes in resynthesized *Brassica napus* and their effect on gene expression and phenotype. The Plant Cell, 19(11), 3403–3417.

Gaeta, R. T., & Pires, J. C. (2010). Homoeologous recombination in allopolyploids: the polyploid ratchet. New Phytologist, 186(1), 18–28.

Gaebelein, R., Schiessl, S. V., Samans, B., Batley, J., & Mason, A. S. (2019). Inherited allelic variants and novel karyotype changes influence fertility and genome stability in *Brassica* allohexaploids. New Phytologist, 223(2), 965–978.

Gonzalo, A., Lucas, M. O., Charpentier, C., Sandmann, G., Lloyd, A., & Jenczewski, E. (2019). Reducing MSH4 copy number prevents meiotic crossovers between non-homologous chromosomes in *Brassica napus*. Nature Communications, 10(1), 1–9.

Gout, J-F., Hao, Y., Johri, P., Arnaiz, O., Doak, T.G., Bhullar, S., Couloux, A., Guerin, F., Malinsky, S., Potekhin, A., Sawka, N., Sperling, L., Labadie, K., Meyer, E., Duharcourt, S., & Lynch, M. (2023). Dynamics of gene loss following ancient whole-genome duplication in the cryptic *Paramecium* complex. Molecular Biology and Evolution, 40(5), msad107.

Hao, Y., Fleming, J., Petterson, J., Lyons, E., Edger, P.P, Pires, J.C., Thorne, J.L., and Conant, G. (2022). Convergent evolution of polyploid genomes from across the eukaryotic tree of life. G3 12(6): jkac094.

Hao, Y., Mabry, M, Edger, P.P, Jin, L, Chuafuang X, VanBuren, R., Colle, M., An, H., Abrahams, R.S., Qi, X, Sankoff, D., Barker, M.S., Lyons, E., Pires, J.C., and Conant, G.C. (2021). The contributions from the progenitor genomes of the mesopolypoid Brassiceae are evolutionarily distinct but functionally compatible. Genome Research 31: 799–810.

He, Z., Wang, L., Harper, A. L., Havlickova, L., Pradhan, A. K., Parkin, I. A., & Bancroft, I. (2017). Extensive homoeologous genome exchanges in allopolyploid crops revealed by mRNA seq-based visualization. Plant Biotechnology Journal, 15(5), 594–604.

Higgins, E.E., Howell, E.C., Armstrong, S.J., & Parkin, I.A.P. (2021) A major quantitative trait locus on chromosome A9, BnaPh1, controls homoeologous recombination in Brassica napus. New Phytologist, 229(6), 3281–3293.

Hou, J., Shi, X., Chen, C., Islam, M. S., Johnson, A. F., Kanno, T., … & Birchler, J. A. (2018). Global impacts of chromosomal imbalance on gene expression in Arabidopsis and other taxa. Proceedings of the National Academy of Sciences, 115(48), E11321–E11330.

Hurgobin, B., Golicz, A. A., Bayer, P. E., Chan, C. K. K., Tirnaz, S., Dolatabadian, A., … & Edwards, D. (2018). Homoeologous exchange is a major cause of gene presence/absence variation in the amphidiploid *Brassica napus*. Plant Biotechnology Journal, 16(7), 1265–1274.

Jenczeswki, E., Eber, F., Grimaud, A., Huet, S., Lucas, M.O., Monod, H., & Chevre, A.M. (2003). *PrBn*, a major gene controlling homeologous pairing in oilseed rape (Brassica napus) haploids. Genetics, 164(2), 645–653.

Kassambara, Alboukadel (2020). ggpubr: ‘ggplot2’ Based Publication Ready Plots. R package version 0.4.0. https://CRAN.R-project.org/package=ggpubr

Kwon, C-T, Tang, L., Want, X., Gentile, I., Hendelman, A., Robitaille, G., Van Eck., J., Xu, C., & Lippman, Z.B. (2022). Dynamic evolution of small signalling peptide compensation in plant stem cell control. Nature Plants, 8, 346–355.

Leal-Bertioli, S.C.M., Godoy, I.J., Santos, J.F., Doyle, J.J., Guimaraes, P.M, Abernathy, B.L., Jackson, S.A., Moretzsohn, M.C., & Bertioli, D.J. (2018). Segmental allopolyploidy in action: Increasing diversity through polyploid hybridization and homoeologous recombination. American Journal of Botany, 105(6), 1053–1066.

Li, Z., Defoort, J., Tasdighian, S., Maere, S., Van de Peer, Y., & De Smet, R. (2016). Gene duplicability of core genes is highly consistent across all angiosperms. The Plant Cell, 28(2), 326–344.

Lloyd, A., Blary, A., Charif, D., Charpentier, C., Tran, J., Balzergue, S., … & Jenczewski, E. (2018). Homoeologous exchanges cause extensive dosage-dependent gene expression changes in an allopolyploid crop. New Phytologist, 217(1), 367–377.

Lyons, E., & Freeling, M. (2008). How to usefully compare homologous plant genes and chromosomes as DNA sequences. The Plant Journal, 53(4), 661–673.

Lyons, E., Pedersen, B., Kane, J., Alam, M., Ming, R., Tang, H., … & Freeling, M. (2008). Finding and comparing syntenic regions among Arabidopsis and the outgroups papaya, poplar, and grape: CoGe with rosids. Plant Physiology, 148(4), 1772–1781.

Madlung, A., Tyagi, A. P., Watson, B., Jiang, H., Kagochi, T., Doerge, R. W., … & Comai, L. (2005). Genomic changes in synthetic *Arabidopsis* polyploids. The Plant Journal, 41(2), 221–230.

Maere, S., De Bodt, S., Raes, J., Casneuf, T., Van Montagu, M., Kuiper, M., & Van de Peer, Y. (2005). Modeling gene and genome duplications in eukaryotes. Proceedings of the National Academy of Sciences, 102(15), 5454–5459.

Makino, T., & McLysaght, A. (2010). Ohnologs in the human genome are dosage balanced and frequently associated with disease. Proceedings of the National Academy of Sciences, 107(20), 9270–9274.

Mason, A. S., & Wendel, J. F. (2020). Homoeologous exchanges, segmental allopolyploidy, and polyploid genome evolution. Frontiers in Genetics, 11, 1014.

Mortazavi, A., Williams, B. A., McCue, K., Schaeffer, L., & Wold, B. (2008). Mapping and quantifying mammalian transcriptomes by RNA-Seq. Nature Methods, 5(7), 621–628.

Nicholas, S.D., Le Mignon, G., Eber, F., Coriton, O., Monod, H., Clouet, V., Huteau, V., Lostanlen, A., Delourme, R., Chalhoub, B., Ryder, C.D., Chevre, A.M., & Jenczewski, E. (2007). Homeologous recombination plays a major role in chromosome rearrangements that occur during meiosis of *Brassica napus* haploids. Genetics, 175 (2), 487–503.

Nicholas, S.D., Monod, H., Eber, F., Chevre, A.M., & Jenczewski, E. (2012). Non-random distribution of extensive chromosome rearrangements in *Brassica napus* depends on genome organization. Plant Journal, 70 (4), 691–703.

Orantes-Bonilla M, Makhoul M, Lee H, Chawla HS, Vollrath P, Langstroff A, Sedlazeck FJ, Zou J and Snowdon RJ (2022) Frequent spontaneous structural rearrangements promote rapid genome diversification in a *Brassica napus* F1 generation. Frontiers in Plant Science 13:1057953. doi: 10.3389/fpls.2022.1057953

Osborn, T.C., Butrulle, D.V., Sharpe, A.G., Pickering, K.J., Parkin, I.A.P., & Lydiate, D.J. (2003). Detection and effects of a homeologous reciprocal transposition in *Brassica napus*. Genetics, 165(3), 1596–1577.

Parey, E., Louis, A, Montfort, J., Guiguen, Y., Roest Crollius, H., & erthelot. C. (2022). An atlas of fish genome evolution reveals delayed rediploidization following the teleost whole-genome duplication. Genome Research. 32(9), 1685–1697.

Paterson, A. H., Chapman, B. A., Kissinger, J. C., Bowers, J. E., Feltus, F. A., & Estill, J. C. (2006). Many gene and domain families have convergent fates following independent whole-genome duplication events in *Arabidopsis, Oryza, Saccharomyces* and *Tetraodon*. Trends in Genetics, 22(11), 597–602.

Pelé, A., Rousseau-Gueutin, M., & Chèvre, A. M. (2018). Speciation success of polyploid plants closely relates to the regulation of meiotic recombination. Frontiers in Plant Science, 9, 907.

Pires, J. C., Zhao, J., Schranz, M. E., Leon, E. J., Quijada, P. A., Lukens, L. N., & Osborn, T. C. (2004). Flowering time divergence and genomic rearrangements in resynthesized Brassica polyploids (Brassicaceae). Biological Journal of the Linnean Society, 82(4), 675–688.

Qi, X, An, H., Hall, T.E., Di, C., Blischak, P.D., Pires, J.C., and Barker, M.S. (2021). Genes derived from ancient polyploidy have higher genetic diversity and are associated with domestication in *Brassica rapa*. New Phytologist 230: 372–386.

R Core Team (2020). R: A language and environment for statistical computing. R Foundation for Statistical Computing, Vienna, Austria. URL https://www.R-project.org/.

Renny-Byfield, S., Gong, L., Gallagher, J. P., & Wendel, J. F. (2015). Persistence of subgenomes in paleopolyploid cotton after 60 my of evolution. Molecular Biology and Evolution, 32(4), 1063–1071.

Renny-Byfield, S., Rodgers-Melnick, E., & Ross-Ibarra, J. (2017). Gene fractionation and function in the ancient subgenomes of maize. Molecular Biology and Evolution, 34(8), 1825–1832.

Rodriguez-Leal, D., Lemmon, Z.H., Man, J., Bartlett, M.E., & Lippman, Z.B. (2017). Engineering quantitative trait variation for crop improvement by genome editing. Cell, 171(2), 470–480.

Rousseau-Guetin, M., Morice, J., Coriton, O., Huteau, V., Trotoux, G., N’egre, S., Falentin, C., Deniot, G., Gilet, M., Eber F.,…& Chevre, A.M. (2017). The impact of open pollination on the structural evolutionary dynamics, meiotic behavior, and fertility of resynthesized allotetraploid *Brassica napus L*. G3: Genes, Genomes, Genetics, 7(2), 705–717.

Salman-Minkov, A., Sabath, N., & Mayrose, I. (2016) Whole-genome duplication as a key factor in crop domestication. Nature Plants, 2, 16115.

Samans, B., Chalhoub, B., & Snowdon, R. J. (2017). Surviving a genome collision: genomic signatures of allopolyploidization in the recent crop species *Brassica napus*. Plant Genome, 10(3), 1–15.

Schnable, J. C., Springer, N. M., & Freeling, M. (2011). Differentiation of the maize subgenomes by genome dominance and both ancient and ongoing gene loss. Proceedings of the National Academy of Sciences, 108(10), 4069–4074.

Schnable, J. C., Wang, X., Pires, J. C., & Freeling, M. (2012). Escape from preferential retention following repeated whole genome duplications in plants. Frontiers in Plant Science, 3:94. doi: 10.3389/fpls.2012.00094

Sharbrough, Justin Conover, J. L., Gyorfy, M. F., Grover, C. E., Miller, E. R., Wendel, J. F., Sloan, D. B. (2022) Global patterns of subgenome evolution in organelle-targeted genes of six allotetraploid angiosperms, Molecular Biology and Evolution, 39(4), msac074

Sharpe, A.G., Parkin, I.A., Keith, D.J., & Lydiate, D.J. 1995. Frequent nonreciprocal translocations in the amphidiploid genome of oilseed rape (*Brassica napus*). Genome 38 (6), 1112–1121.

Shi, X., Zhang, C., Ko, D. K., & Chen, Z. J. (2015). Genome-wide dosage-dependent and independent regulation contributes to gene expression and evolutionary novelty in plant polyploids. Molecular Biology and Evolution, 32(9), 2351–2366.

Shi, X., Yang, H., Chen, C., Hou, J., Hanson, K. M., Albert, P. S., … & Birchler, J. A. (2021). Genomic imbalance determines positive and negative modulation of gene expression in diploid maize. The Plant Cell, 33(4), 917–939.

Song, K., Lu, P., Tang, K. & Osborn, T.C. (2005). Rapid genome change in synthetic polyploids of *Brassica* and its implications for polyploid evolution. Proceedings of the National Academy of Sciences, 92(17), 7719–7723.

Song, M. J., Potter, B. I., Doyle, J. J., & Coate, J. E. (2020). Gene balance predicts transcriptional responses immediately following ploidy change in *Arabidopsis thaliana*. The Plant Cell, 32(5), 1434–1448.

Stebbins, G.L. (1947). Types of polyploids: Their classification and significance. Advances in Genetics 1: 403–429.

Stein, A., Coriton, O., RousseaulJGueutin, M., Samans, B., Schiessl, S. V., Obermeier, C., … & Snowdon, R. J. (2017). Mapping of homoeologous chromosome exchanges influencing quantitative trait variation in *Brassica napus*. Plant Biotechnology Journal, 15(11), 1478–1489.

Szadkowski, E., Eber, F., Huteau, V., Lode, M., Huneau, C., Belcram, H., … & Chèvre, A. M. (2010). The first meiosis of resynthesized *Brassica napus*, a genome blender. New Phytologist, 186(1), 102–112.

Tasdighian, S., Van Bel, M., Li, Z., Van de Peer, Y., Carretero-Paulet, L., & Maere, S. (2017). Reciprocally retained genes in the angiosperm lineage show the hallmarks of dosage balance sensitivity. The Plant Cell, 29(11), 2766–2785.

Thomas, B. C., Pedersen, B., & Freeling, M. (2006). Following tetraploidy in an Arabidopsis ancestor, genes were removed preferentially from one homeolog leaving clusters enriched in dose-sensitive genes. Genome Research, 16(7), 934–946.

Thompson, A., Zakon, H. H., & Kirkpatrick, M. (2016). Compensatory drift and the evolutionary dynamics of dosage-sensitive duplicate genes. Genetics, 202(2), 765–774.

Turner-Hissong, S.D., Mabry, M.E., Beissinger, T.M., Ross-Ibarra, J. and Pires, J.C. (2020). Evolutionary insights into plant breeding. Current Opinion in Plant Biology 54: 93–100.

Vicient, C. M., & Casacuberta, J. M. (2017). Impact of transposable elements on polyploid plant genomes. Annals of Botany. 120(2), 195–207

Wendel, J. F., Lisch, D., Hu, G., & Mason, A. S. (2018). The long and short of doubling down: polyploidy, epigenetics, and the temporal dynamics of genome fractionation. Current Opinion in Genetics & Development, 49, 1–7.

Woodhouse, M. R., Schnable, J. C., Pedersen, B. S., Lyons, E., Lisch, D., Subramaniam, S., & Freeling, M. (2010). Following tetraploidy in maize, a short deletion mechanism removed genes preferentially from one of the two homeologs. PLoS biology, 8(6), e1000409.

Woodhouse, M. R., Cheng, F., Pires, J. C., Lisch, D., Freeling, M., & Wang, X. (2014). Origin, inheritance, and gene regulatory consequences of genome dominance in polyploids. Proceedings of the National Academy of Sciences, 111(14), 5283–5288.

Wu, Y., Lin, F., Zhou, Y., Wang, J., Sun, S., Wang, B., … & Liu, B. (2021). Genomic mosaicism due to homoeologous exchange generates extensive phenotypic diversity in nascent allopolyploids. National Science Review, 8(5), nwaa277.

Xiong, Z., Gaeta, R. T., & Pires, J. C. (2011). Homoeologous shuffling and chromosome compensation maintain genome balance in resynthesized allopolyploid *Brassica napus*. Proceedings of the National Academy of Sciences, 108(19), 7908–7913.

Xiong, Z, Gaeta, R.T., Edger, P.P, Cao, Y, Zhao, K, Zhang, S, & Pires, J.C. (2021). Chromosome inheritance and meiotic stability in allopolyploid *Brassica napus*. G3: Genes, Genomes, Genetics 11: jkaa011.

Yang, H., Shi, X., Chen, C., Hou, J., Ji, T., Cheng, J., & Birchler, J. A. (2021). Predominantly inverse modulation of gene expression in genomically unbalanced disomic haploid maize. The Plant Cell, 33(4), 901–916.

Zhang, Z., Gou, X., Xun, H., Bian, Y., Ma, X., Li, J., … & Levy, A. A. (2020). Homoeologous exchanges occur through intragenic recombination generating novel transcripts and proteins in wheat and other polyploids. Proceedings of the National Academy of Sciences, 117(25), 14561–14571.

Zhang, Z., Xun, H., Lv, R., Gou, X., Ma, X., Li, J., Zhao, J., Li N, Gong, L, & Liu, B. (2022). Effects of homoeologous exchange on gene expression and alternative splicing in a newly formed allotetraploid wheat. Plant Journal 111(5), 1267–1282.

