## Supplementary material for "Dosage-sensitivity shapes how genes transcriptionally respond to allopolyploidy and homoeologous exchange in resynthesized Brassica napus": Fig S4

**BnC**

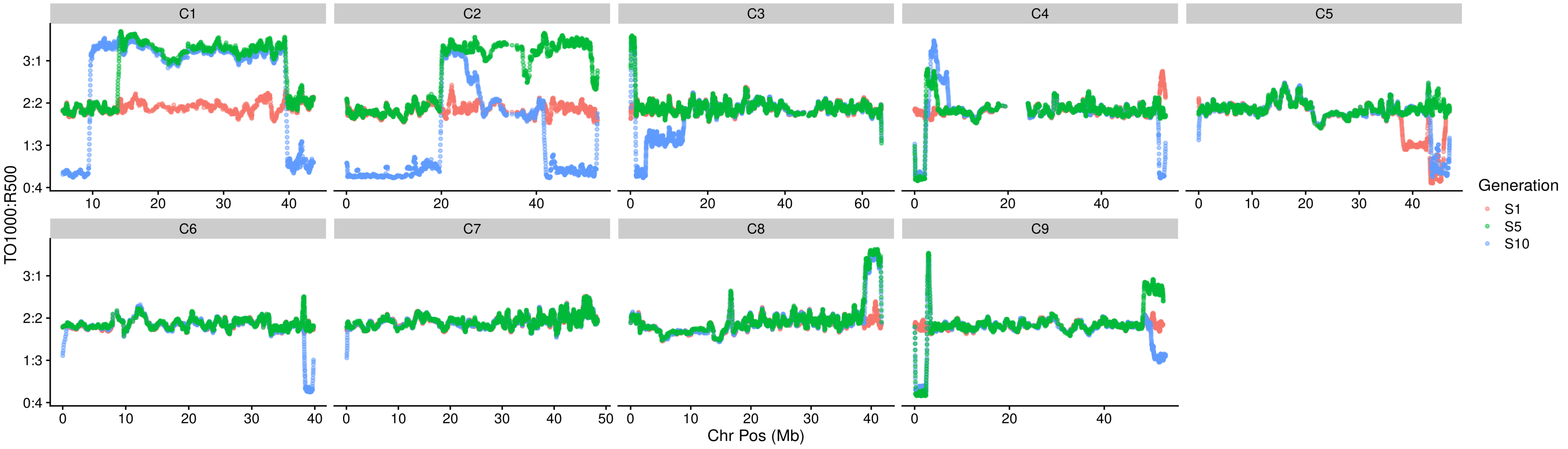


**BnA**

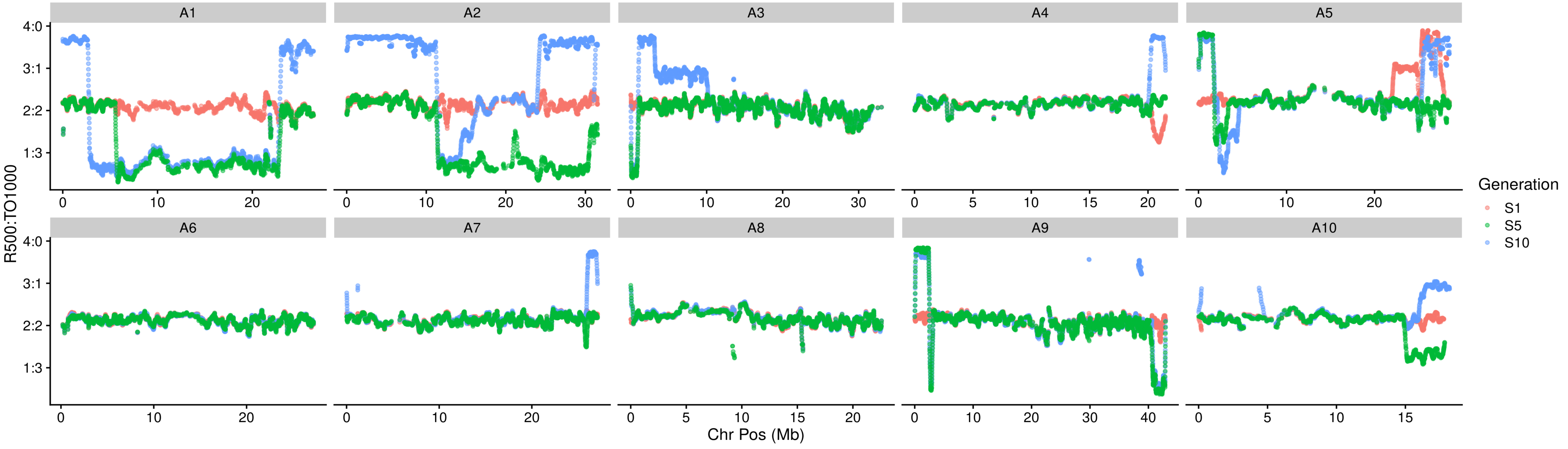


**Fig S4.** Mean read depth ratio of WGS reads for the BnC subgenome (top) and BnA subgenome (bottom) for individual EL400. Values represent the rolling average of the read depth ratio over a 50 gene window. Regions such as chromosomes A1/C1 where long stretches of skewed read depth ratios were taken as evidence of larger genomic changes like aneuploidy or partial duplication/deletion and excluded from subsequent analyses.
