## Supplementary material for "Dosage-sensitivity shapes how genes transcriptionally respond to allopolyploidy and homoeologous exchange in resynthesized Brassica napus": Fig S7

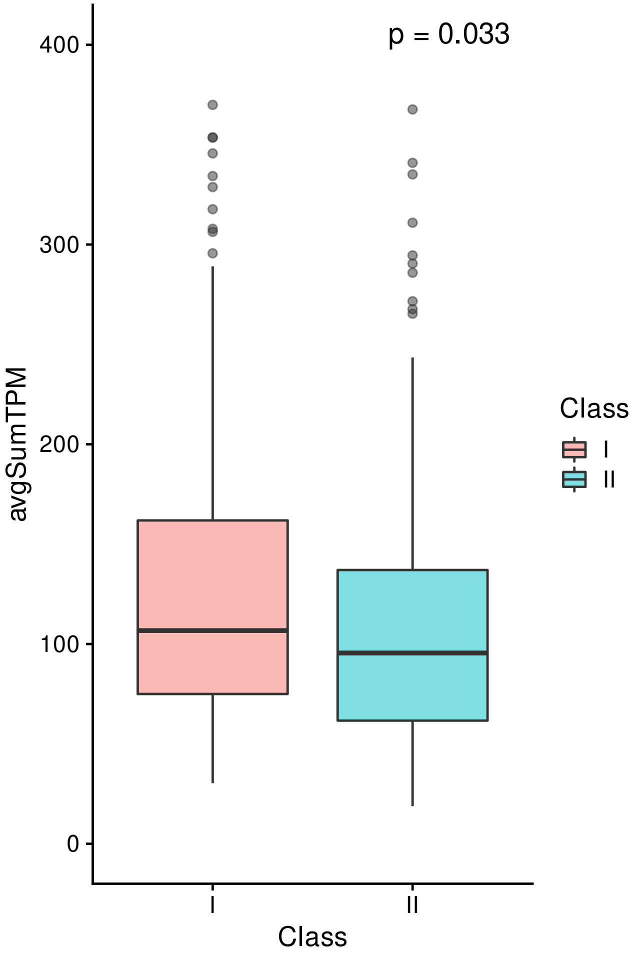


**Fig S7.** Comparison of average TPM of genes from Class I and II GO terms for homoeologous pairs that are affedcted by homoeologous exchange. P-value represents the results of a Kruskal-Wallis test
