## Supplementary material for "Dosage-sensitivity shapes how genes transcriptionally respond to allopolyploidy and homoeologous exchange in resynthesized Brassica napus": Fig S8

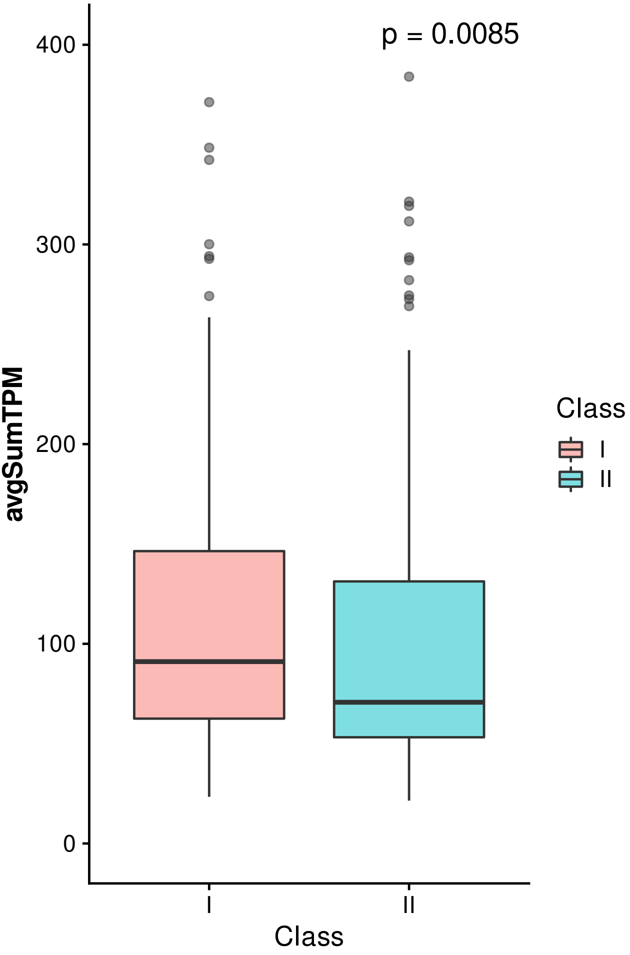


**Fig S8.** Comparison of average TPM of genes from Class I and II GO terms for homoeologous pairs that are at a 2:2 dosage ratio. P-value represents the results of a Kruskal-Wallis test
